# High-resolution structure of mouse radial spoke and its in-situ structure in ependymal cilia revealed by cryo-EM and cryo-ET

**DOI:** 10.1101/2023.06.02.543457

**Authors:** Xueming Meng, Cong Xu, Jiawei Li, Benhua Qiu, Jiajun Luo, Qin Hong, Yujie Tong, Chuyu Fang, Yanyan Feng, Rui Ma, Xiangyi Shi, Cheng Lin, Chen Pan, Xueliang Zhu, Xiumin Yan, Yao Cong

**Affiliations:** State Key Laboratory of Molecular Biology, Shanghai Institute of Biochemistry and Cell Biology, Center for Excellence in Molecular Cell Science, Chinese Academy of Sciences, Shanghai, China; University of Chinese Academy of Sciences; State Key Laboratory of Cell Biology, Shanghai Institute of Biochemistry and Cell Biology, Center for Excellence in Molecular Cell Science, Chinese Academy of Sciences, Shanghai, China; University of Chinese Academy of Sciences; Ministry of Education-Shanghai Key Laboratory of Children’s Environmental Health, Institute of Early Life Health, Xinhua Hospital, Shanghai Jiao Tong University School of Medicine, Shanghai 200092, China; Shanghai Nanoport, Thermofisher Scientific, Shanghai, China; National Facility for Protein Science in Shanghai, Shanghai Advanced Research Institute, Chinese Academy of Sciences, Shanghai 201210, China; Key Laboratory of Systems Health Science of Zhejiang Province, School of Life Science, Hangzhou Institute for Advanced Study, University of Chinese Academy of Sciences, Hangzhou, China

## Abstract

Radial spokes (RS) transmit mechanochemical signals between the central pair (CP) and axonemal dynein arms to coordinate ciliary motility. However, the high-resolution structure of mammalian RS remains missing. Here, we present the high-resolution cryo-EM structure of mouse RS head-neck complex in both monomer and dimer forms and reveal the intrinsic dynamics of the dimer. We also map the genetic mutations related to primary ciliary dyskinesia and asthenospermia on the head-neck complex. Moreover, we present the unprecedented cryo-ET and sub-tomogram averaging map of mouse ependymal cilia, whose beating creates unidirectional cerebrospinal fluid flow, and build the models for RS1-3, IDAs, and N-DRC. Strikingly, our cryo-ET map reveals the lack of IDA-b/c/e and the absence of tektin filaments within the A-tubule of doublet microtubules in ependymal cilia compared with mammalian respiratory cilia and sperm flagella. This tissue-specific feature may represent an evolutionary choice driven by the functional requirements on ependymal cilia. Our findings reveal the RS head-neck assembly mechanism, shed light on the coordinated rigid and elastic RS-CP interaction modes beneficial for the regulation of asymmetric ciliary beating, and also facilitate understandings on the etiology of ciliary dyskinesia-related ciliopathies.

## Introduction

Motile cilia or flagella are hair-like organelles functioning mainly as “paddles” to drive rapid cell movement or extracellular fluid flow. In vertebrates, motile cilia reside on the epithelial cell surface of various tissues, including the ependyma, trachea, and fallopian tubes, whereas flagella are solely present in sperm. Dysfunctions in motile cilia and/or flagella lead to primary ciliary dyskinesia (PCD), which is characterized by chronic respiratory disease, organ inversions, hydrocephalus, and infertility^1,2^. For instance, ependymal multicilia, which line the surface of brain ventricle walls, create a unidirectional cerebrospinal fluid (CSF) flow through their coordinated back-and-forth beating. As CSF is rich in neuropeptides, its orderly flow is critical for nourishing the central nervous system and maintaining proper body axis. Dyskinesia of ependymal multicilia leads to the obstruction of CSF flow, resulting in hydrocephalus and idiopathic scoliosis^3–5^.

The axoneme of motile cilia and flagella is generally composed of nine peripheral doublet microtubules (DMTs) surrounding a central pair (CP) of MTs (the “9+2” axoneme). Along the DMTs, seven inner dynein arms (IDAs), four outer dynein arms (ODAs), and three radial spokes (RSs) are arranged in 96-nm repeats to generate ciliary movement together with the CP^6,7^. The RS is a T-shaped protein complex with an orthogonal head pointing toward the CP and a stalk anchored on each A-tubule of the DMTs^7,8^. It acts as the mechanochemical transducer between the CP and axonemal dynein arms to regulate flagellar/ciliary motility^9–12^. The RS is a macromolecular machinery consisting of more than twenty subunits. The flagella of *Chlamydomonas*, a widely used protist model organism, contain two full-size RSs (RS1 and RS2) and a truncated RS3 with only the stalk in each 96-nm repeat unit of the axoneme. In contrast, motile cilia and flagella of most other protists, such as *Tetrahymena thermophile*, and metazoa possess triplet RSs (RS1 to RS3)^7,8^. Genetic mutations in the RS head-neck components also cause PCD^13–18^.

During the past years, using cryo-electron tomography (cryo-ET) and sub-tomogram averaging, significant progress has been made in understanding the structures of RS in various organisms, including protists such as *Chlamydomonas*, *Tetrahymena*, *Choanoflagellate*, and *Trypanosoma*, as well as metazoa such as human or bovine respiratory cilia and sea urchin, mouse, or human sperm flagella^7,8,16, 19–22^. These efforts have led to a better understanding of morphologies, compositions, and variations of the RSs. We have previously determined the high-resolution cryo-EM structure of the mammalian RS head monomer^23^. Meanwhile, high-resolution cryo-EM structures of *Chlamydomonas* RS head and head-neck have been reported^24,25^. These studies have revealed marked differences in morphology and composition between metazoan RSs and *Chlamydomonas* ones^7,8,19, 23–31^. Interestingly, even within mammals, ciliary and flagellar RSs differ in morphology and composition^21–23^.

Here, we reconstituted a mouse RS head-neck complex and determined its unprecedented high-resolution cryo-EM structures in both monomer and dimer forms to a resolution up to 3.14 Å. Our study revealed that the intrinsic dynamics of the RS head-neck dimer, combined with the proposed RS-CP contact modes, contribute to the elegant regulation mechanism of ciliary motility. We also provided potential etiology of RS head-neck gene mutations linked to PCD and asthenospermia^13,32,33^. Furthermore, we present the first *in situ* axonemal structure of mouse ependymal cilia, including multi-scale models for RS1, RS2, RS3, IDAs, and N-DRC based on our cryo-EM and cryo-ET structures and AlphaFold2 predicted models. Altogether, our results reveal the assembly mechanism of mammalian RS head-neck for RS1 and RS2, the mode of mammalian RS-CP interactions, and the structure-function relationships of ciliary and flagellar axonemes.

## Results

### Reconstitution of a stable mouse heteroheptameric RS head-neck complex

In a previous study, we successfully reconstituted the mouse RS head complex, which consists of Rsph1-Rsph4a-Rsph9-Rsph3b subunits and exhibits a two-fold symmetric brake pad-shaped structure^23^. On this basis, we aimed to assemble and gain insights into a more complete mouse RS head-neck complex. To determine whether the RS head and neck subunits can form a stable complex, we co-expressed Flag-tagged Rsph16 and *Strep*-Tag II-tagged Rsph1, together with Rsph3b, -4a, -9, -10b, -23, and -2 in HEK-293F cells, and then purified the complex by a two-step affinity purification strategy. SDS-PAGE followed by Coomassie brilliant blue staining indicated that a complex composed of Rsph1-Rsph4a-Rsph9-Rsph3b-Rsph23-Rsph2-Rsph16 was readily detected (Fig. 1A). The complex was further confirmed by glycerol-gradient ultracentrifugation and mass spectrometry analysis (Fig. S1A-B).

**Fig. 1.**
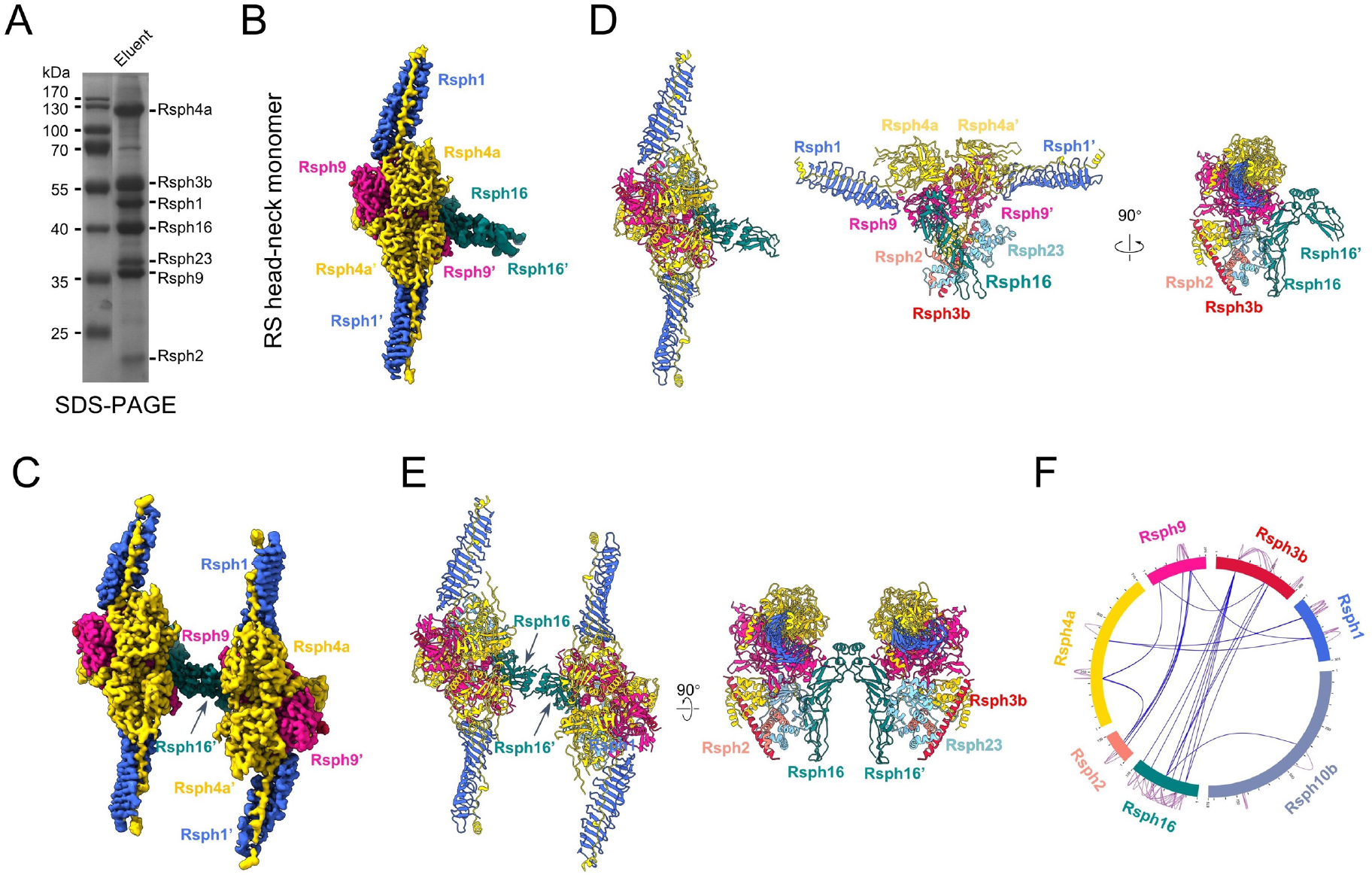
Cryo-EM structures of mouse RS head-neck complex. (A) Coomassie-stained SDS– PAGE of the RS head-neck complex. (B, C) Cryo-EM map of the RS head-neck monomer (B) and dimer (C), with subunits shown in distinct colors. This color schema is followed throughout. (D, E) Atomic model for the RS head-neck monomer (D) and dimer (E). (F) XL-MS analysis of the RS head-neck complex. Identified cross-linked inter-subunit contacts are shown as blue lines, and intra-subunit contacts as purple lines. We used best E-value (1.00E-02) and spec count of at least 2 as the threshold to remove extra XL-MS data with lower confidence.

### The RS head-neck complex exists as both monomer and dimer

Our cryo-EM analysis on the assembled mouse RS head-neck complex revealed the presence of both monomer and dimer (Fig. S1C-D). We determined their structures at a resolution of 3.28 Å and 3.57 Å, respectively, with the head core region focus-refined to a resolution of 3.14 Å (Fig. 1B-C, S2, S3A-C, Table S1). In this dataset, we also detected a less populated RS head structure (Fig. S2) that resembles the one obtained in our previous study^23^ (Fig. S3D). We then built atomic models for the RS head-neck monomer and dimer based on our previous mouse RS head structure^23^ and AlphaFold2 predicted models for the neck subunits, with constraints of the current maps (Fig. 1D-E). The model fits in the corresponding map very well (Fig. S3E, G, Movie S1).

In the head-neck monomer structure, the head portion consists of a compact body and two extending Rsph1 arms, made up of two copies of Rsph1/Rsph4a/Rsph9 (Fig. 1B, D), resembling the structure of the RS head in our previous study^23^. Compared with the *Chlamydomonas* head, which consists of RSP4/RSP6/RSP9/RSP9’/RSP1/RSP10/RSP5/RSP2 ^24,25,30^, the mammalian RS head appears to be much more reduced in composition and morphology (Fig. S4A-B). In our previous study^23^, we were unable to resolve the structure of Rsph3b. However, in the current RS head-neck complex, Rsph3b is attached to one side of the head and wrapped around by the previously missing N-terminal Dpy-30 motif of the head subunits Rsph4a and Rsph4a’ (Fig. 1D), as well as the Dpy-30 motif of the neck subunits Rsph2 and Rsph23. This leads to a fluent connection between the head and the neck, which is in line with that observed in *Chlamydomonas*^24,25^. This RS head-neck associates laterally with one side of an arch bridge composed of two Rsph16 subunits to form the RS head-neck monomer (Fig. 1B, D). In contrast, the RS head-neck dimer consists of two copies of the RS head-neck connected in between through the arch bridge of Rsph16-Rsph16’ (Fig. 1C, E).

We next performed XL-MS analysis on the RS head-neck complex (Fig. 1F, Table S3), and the results overall support the subunit spatial arrangement determined by the cryo-EM study. Notably, both our mass spectrometric and XL-MS results indicated the presence of Rsph10b in the purified complex (Fig. 1F, S1B). Unlike the extensive interplays among the other subunits, however, Rsph10b only exhibits limited association with Rsph16 in the XL-MS results (Fig. 1F), suggesting that Rsph10b may not be a standard component of RS1 and RS2’s head-neck complex.

### Subunit interaction network of the RS head-neck complex

Compared with our previously reported subunit interaction network within the RS head^23^, the current RS head-neck structure disclosed additional interactions in the head-neck interface and within the neck (Fig. 2A, Table S4). Firstly, the Rsph4a N-terminal extension wraps around the head and involves intensive interactions with the head subunit Rsph9 through rich H-bonds/salt-bridges and the neck subunit Rsph3b (Fig. 2A, 1-2). Secondly, the Dpy-30 motifs from not only Rsph4a and Rsph4a’ but also Rsph2 and Rsph23 wrap around the long α-helix of Rsph3b (Fig. 2A, 2, S4C), so that Rsph3b intimately engages the head and the neck subunits. Thirdly, one RS head-neck complex attaches to each side of the arch bridge formed by the Rsph16 dimer. Close inspection of this area revealed that the RS head-neck is connected with the arch bridge through an intense H-bond/salt-bridge interaction network between Rsph9 and Rsph16 (Fig. 2A, 3) and H-bond interactions between Rsph23 and Rsph16 (Fig. 2A, 4). Finally, in the arch bridge, the Rsph16-Rsph16’ interaction appears to occur in two areas, involving electrostatic interactions between their C-terminal α-helix (Fig. 2A, 5) and hydrophobic interactions between a part of the DNAJ-C domain (Fig. 2A, 6, S4C).

**Fig. 2.**
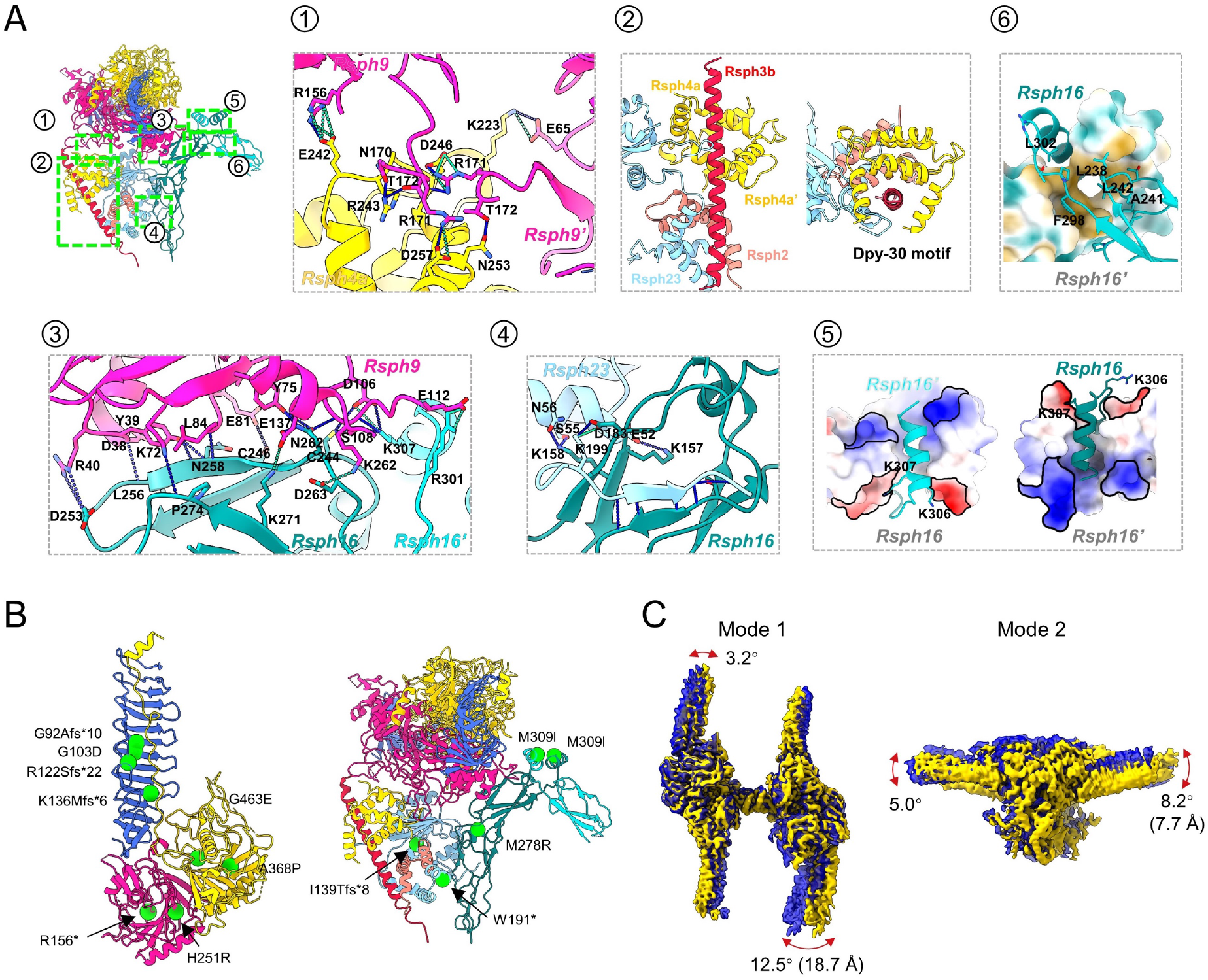
Subunit interaction networks and mapping of the PCD/asthenospermia-related mutations on the RS head-neck complex, as well as the intrinsic dynamics of the head-neck dimer. (A) The interaction networks of Rsph9-Rsph4a (1), between Rsph3b and the Dpy-30 domains of Rsph4a/Rsph4a’/Rsph2/Rsph23 (2), of Rsph9-Rsph16 (3), and Rsph23-Rsph16 (4). The electrostatic interaction between the C-terminal a-helix of Rsph16 and Rsph16’ (5), and the hydrophobic interaction between the Rsph16 and Rsph16’ (6). H-bonds are drawn in blue dash line and salt bridges in green dash line. (B) Known PCD/asthenospermia disease-causing mutations mapped onto the model of the RS head-neck complex. Each green ball represents a mutation associated with PCD/asthenospermia disease. (C) Two representative modes of motion of the RS head-neck dimer. The angular range and direction as well as the extreme distance of the motion are displayed on the overlaid two extreme maps in the motion (in blue and yellow).

**Fig. 3.**
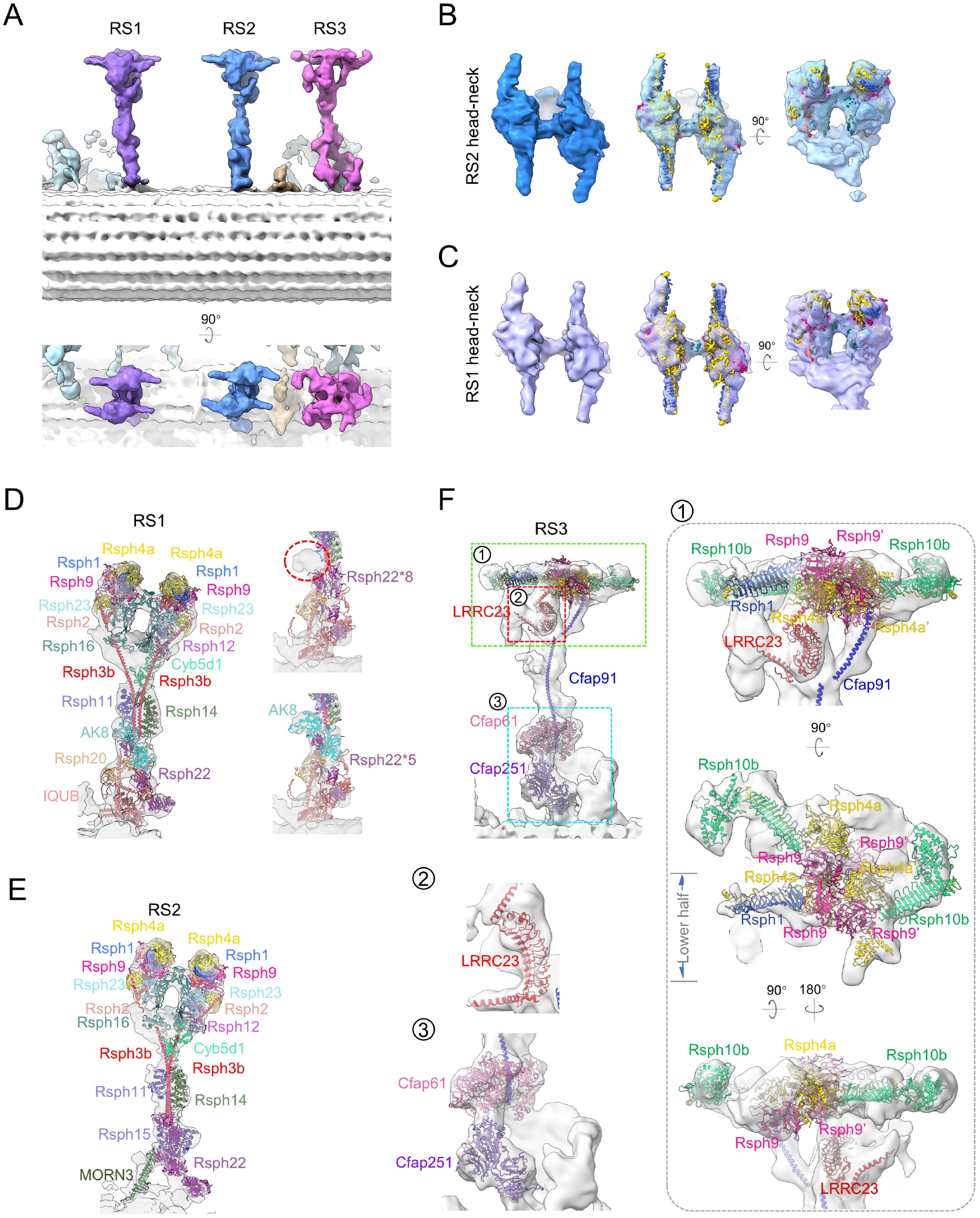
The *in situ* cryo-ET structure of the mouse ependymal cilia axoneme and the proposed multi-scale models for RS1/RS2/RS3. (A) Different views of the cryo-ET map of the 96-nm axonemal repeat of mouse ependymal cilia, with RS1/RS2/RS3 shown in distinct color. (B, C) Fitting of our cryo-EM map of the RS head-neck dimer (in color) into the corresponding head-neck region of the *in situ* cryo-ET map of RS2 (B, blue) or RS1 (C, purple), indicating a perfect matching between them. (D) A proposed model of mammalian RS1, with our RS head-neck dimer model and AlphaFold2 predict models for the stalk/base fit into the cryo-ET map of RS1. We also placed an AK8 subunit in the stalk region of RS1, replacing several Rsph22 (right panels). (E) A proposed model of mammalian RS2. (F) A proposed model for the mammalian RS3. The RS3 head-neck is displayed in different views in (1).

**Fig. 4.**
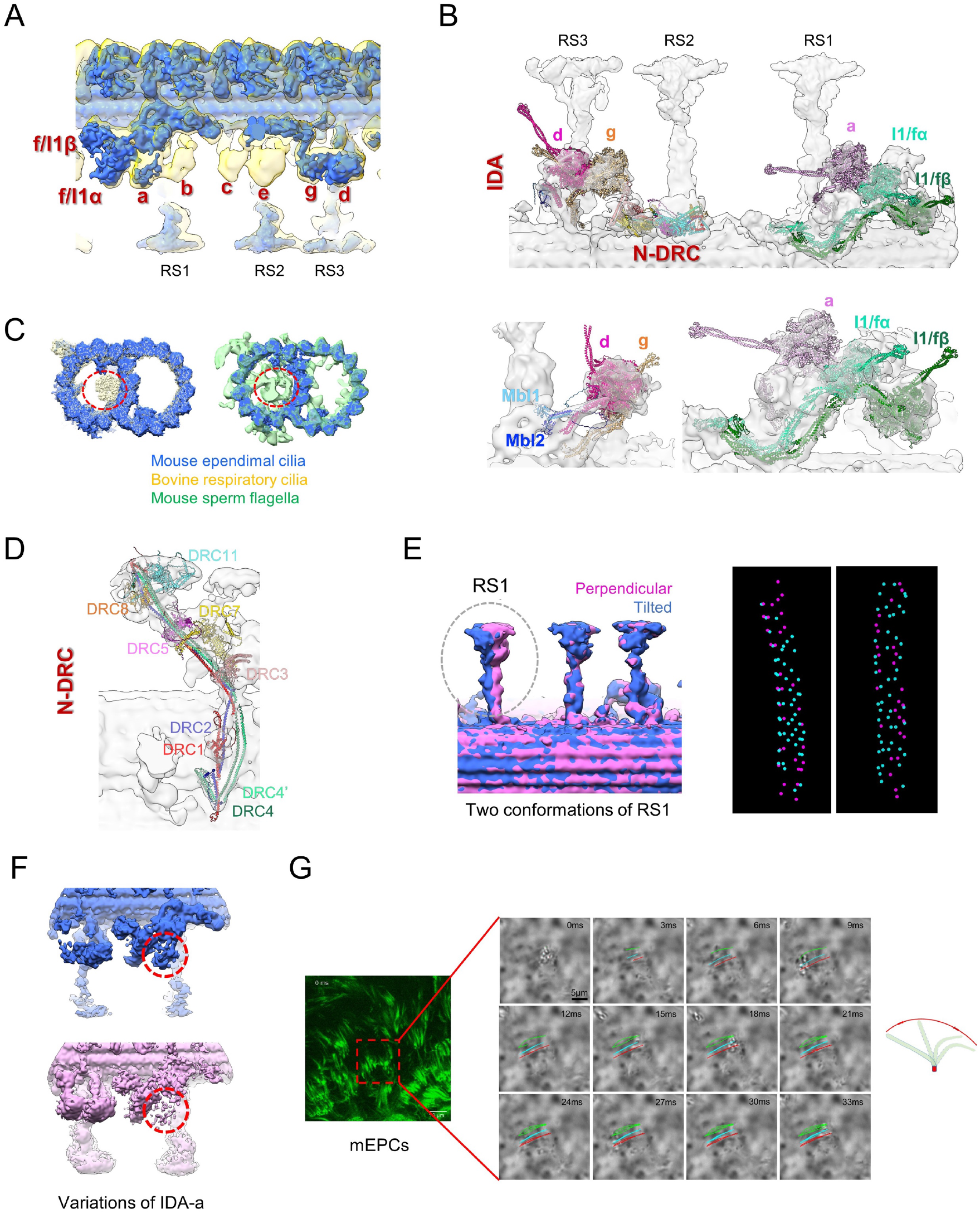
Reduced IDAs and MIPs in DMT of mouse ependymal axoneme revealed by cryo-ET. (A) Overlaid cryo-ET maps of mouse ependymal cilia axoneme (royal blue) with human respiratory ciliary axoneme (transparent yellow, EMD-5950) revealed that IDA-a/b/c are absent in mouse ependymal cilia. (B) Proposed models for the IDAs of mouse ependymal cilia. (C) Cross-section view of the overlaid cryo-ET maps of the mouse ependymal cilia DMT 48-nm repeat (royal blue) vs. the bovine respiratory DMT (transparent kaki, EMD-24664) or mouse sperm flagella (transparent green, EMD-27455), revealed greatly reduced MIPs (e.g., missing tektins) within the A-tubule of ependymal cilia DMT (indicated by dotted red circles). (D) Proposed model fits into the N-DRC cryo-ET map of mouse ependymal cilia. (E) 3D classification revealed two *in situ* conformations of RS1 (left) and the *in situ* distribution of the two conformations in two representative tomograms (right). The perpendicular RS1 conformation is shown in magenta, and the tilted one in blue. (F) Focused 3D classification revealed two *in situ* IDA-a states (indicated by dotted red circle), in one of which the IDA-a appears nearly missing (lower panel). (G) The beating pattern of mouse ependymal cilia.

**Fig. 5.**
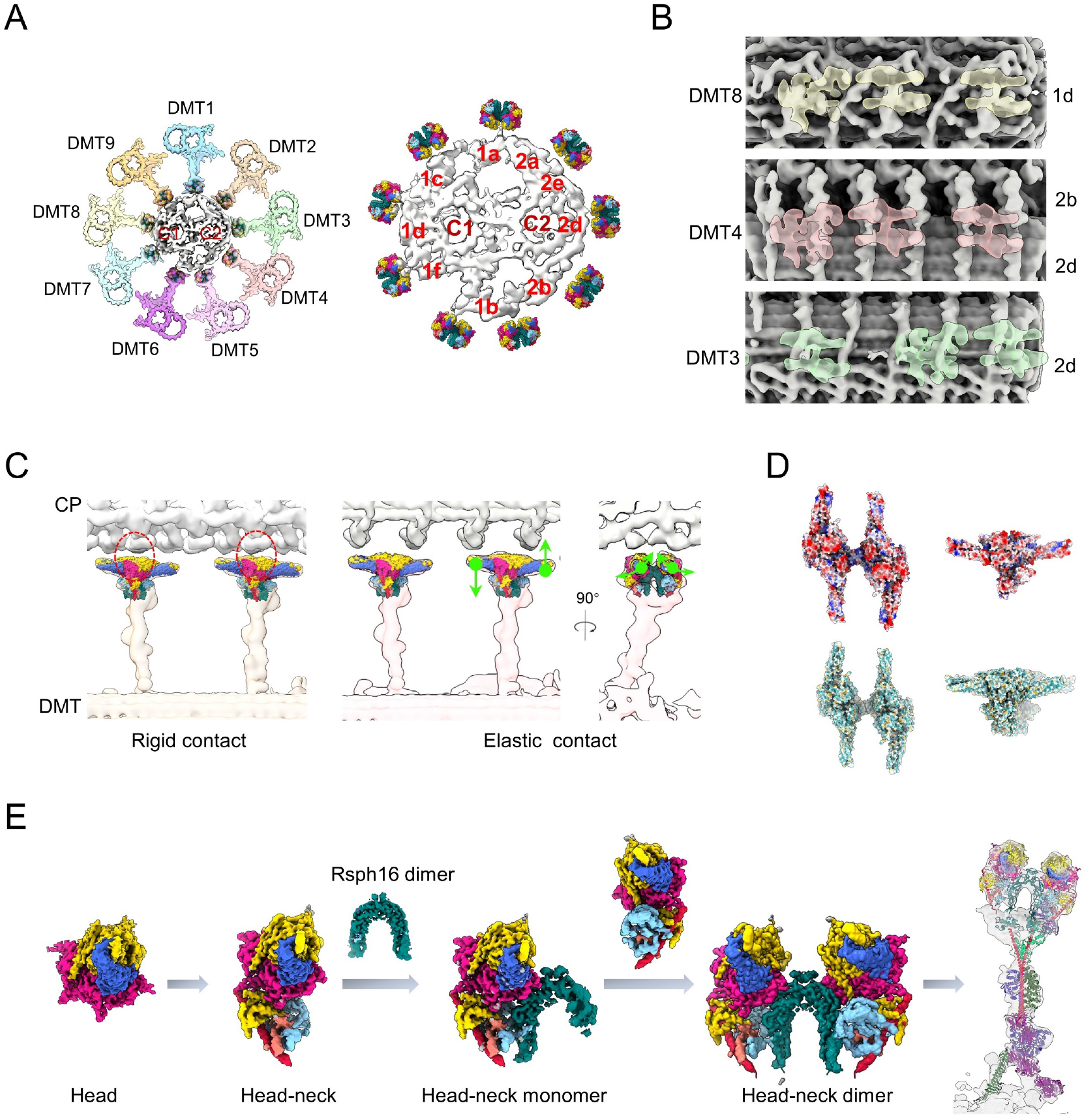
Proposed RS-CP interaction patterns and the assembly mechanism for mammalian RS head-neck complex. (A) Coordination of the RS head-neck complex-fitted DMT of our mouse ependymal cilia and the recent cryo-ET map of mouse sperm CP (EMD-27445) into the framework of a 9+2 axoneme by matching remaining RS densities of the CP. (B) Longitudinal views of locations of the RSs (transparent density) relative to the indicated periodic CP projections (in grey). The representative RSs from DMT3/4/8, better matched with the remaining RS densities of the CP, are shown here. (C) Proposed RS-CP interaction patterns, in coordination with the intrinsic dynamics of RS head-neck complex. Left: rigid contact. Right: seesaw-like alternating up-and-down motions (left) and open-and-close motion (right). (D) The surface properties of the RS head-neck dimer. (E) The proposed assembly mechanism for the mammalian RS head-neck complex of RS1/RS2.

### Mapping of the PCD**/**asthenospermia-related mutations on the RS head-neck complex

Since many pathogenic mutations of PCD or asthenospermia patients on the RS head-neck subunits have been documented (Table S5)^15,17,34^, our atomic model of the head-neck complex enabled us to map these mutations on this structure (Fig. 2B). Frameshift mutations such as G92Afs*10, R122Sfs*22, and K136Mfs*6 in the MORN motif of Rsph1^15^ truncate the protein and might also disrupt its interaction with Rsph4a, thus disturbing the arm formation of the RS head (Fig. 2B). Similarly, the W191* mutation of Rsph23^35^ might disrupt the head-neck assembly by truncating the protein (Fig. 2B). The I139Yfs*8 mutation of Rsph23 truncate its C-terminus^35,36^, which is responsible for its binding to Rsph16 to stabilize the head-neck complex, and may therefore induce instability of the complex.

Missense mutations in Rsph16, such as M278R and M309I, have been reported to be related to male fertility in asthenozoospermia^13,32^. The M278R mutation changes the hydrophobic Met into a hydrophilic Arg with a longer sidechain, which may result in a potential clash and repulsion with the adjacent R225, which could be propagated to R189 located on the other side of R225 through H-bond networks (Fig. S4D). R189 resides in the loop involved in the interaction of Rsph16 with the C-terminal tail of Rsph23, thus eventually disturbing the interaction of Rsph16 with Rsph23 and weakening the stability of the head-neck complex. The M309I mutation is located at the Rsph16 dimerization interface formed by the C-terminal α-helix (Fig. 2B). The mutation of Met into Ile may disturb the formation of the arch bridge and impair the stability or even formation of the head-neck dimer.

### Conformational dynamics of the RS head-neck dimer

Our local resolution analysis of the RS head-neck dimer revealed relatively lower local resolution at the very end of the Rsph1 arm and the neck (Fig. S3B), indicating plasticity in these regions. To further examine the intrinsic dynamics of the complex, we performed 3D flexible refinement on the head-neck dimer in cryoSPARC^37^ and found considerable relative movements between the two monomers (Fig. 2C). Specifically, Mode 1 obtained through this analysis mainly describes an open-and-close movement between the Rsph1 arms of the two monomers with an angular range of Rsph1-rotation up to 12.5° (Fig. 2C and Movie S2). Mode 2 corresponds more to the seesaw-like alternating up-and-down motions between the two monomers, with an angular range of Rsph1 arm rotation up to 8.2° (Fig. 2C and Movie S3). These intrinsic motions of the head-neck dimer may contribute to the RS-CP interaction and ciliary motion regulation (discussed later).

### The RS head-neck dimer cryo-EM map matches well with the cryo-ET map of RS1 and RS2 in ependymal cilia

To clarify whether the RS head-neck dimer is a *bona fide* one in motile cilia, we determined the *in situ* axonemal structure of ependymal cilia in cultured mouse ependymal cells (mEPCs) using cryo-ET combined with sub-tomogram averaging and achieved a consensus map at the resolution of 24.2 Å (Fig. 3A, S5A, Table S2, Movie 4). We performed further focused refinement on the RS1/RS2/RS3 region to improve structural details (Fig. S5B). We then fitted our high-resolution cryo-EM map of the RS head-neck dimer into the cryo-ET map of RS2 in the head-neck region, and found that they match each other very well (Fig. 3B). This is also the case for the *in situ* RS1 map in the head-neck region (Fig. 3C). Collectively, these data suggested that our recombinant RS head-neck complex represents the endogenous head-neck architecture of both RS1 and RS2, thus further confirming that the mammalian RS head complex is distinct in composition from the *Chlamydomonas* one^7, 23–25,27^.

### Structural models of mammalian RS1/RS2/RS3

To achieve a complete model of the mammalian RS1 and RS2, we first fitted our RS head-neck dimer model into the corresponding head-neck region of our ependymal cilia *in situ* RS1/RS2 maps. We then fitted AlphaFold2 predicted models of several RS stalk/base components into their corresponding locations, based on features of the cryo-ET map and models, as well as previous reports on mammalian^38^ and *Chlamydomonas* RSs^24,27^. For RS1, in addition to the head and neck region, we fitted the models of Cyb5d1, Rsph12, Rsph11, Rsph14, Rsph20, Rsph22, and IQUB in the stalk and base regions (Fig. 3D, Table S6). We also placed an AK8 subunit in the stalk region of RS1, replacing several Rsph22 (Fig. 3D), based on the good fitting between the model and the map as well as our MS analysis (Table S7), suggesting that AK8 exists in the RSs of EPCs. For RS2, in addition to the head-neck region, we also fitted the Cyb5d1, Rsph12, Rsph11, Rsph14, Rsph15, Rsph22, and MORN3 subunits in the stalk and base regions of RS2 (Fig. 3E).

Noteworthily, the composition and the high-resolution structure of RS3 remain largely unknown. To provide structural insights into this unique RS, we managed to fit several potential components into our *in situ* RS3 map based on its structural features and recent reports on the potential composition of mammalian RS3^18,39,40^. For instance, LRRC23 is a potential RS3 component because its absence abolishes the RS3 head in mice and humans^18^. Indeed, the AlphaFold2 predicted model of LRRC23, which displays a characteristic curved feature, matches a bent density underneath the RS3 head very well (Fig. 3F, 2). In line with recent reports suggesting that LRRC23 contacts with Rsph9^18,40^, we observed a head region right above LRRC23 displaying an obvious double hump feature, which matches the core of the RS head (Rsph4a/Rsph9) reasonably well (Fig. 3F, 1). In addition, a previous cryo-ET study on trachea cilia of *Rsph4a*-deficient mice indicated the loss of all triplet RS heads^41^, and another study on respiratory cilia from *Rsph1-* or *Rsph4a*-deficient PCD patients also suggested their involvement in RS3^42^. Collectively, these results indicated that the head subunits Rsph4a/Rsph9/Rsph1 could also exist in RS3. We then fitted a RS head complex, with one Rsph1 arm substituted by a Rsph10b, into the lower half of the RS3 head density (Fig. 3F, 1). In this proposed fitting, the C-terminal globular domain of Rsph10b fits well in a hook-like density (Fig. 3F, 1), and the N-terminal Dyp-30 motif of two Rsph4a subunits held in the wing-like density reasonably well (Fig. 3F, 1). Moreover, in the other half of the RS3 head, a head core containing two Rsph9 and one Rsph4a could be fitted into a hump-like region (Fig. 3F, 1), and the long loop of Rsph4a could snugly fit into the MORN motif of another Rsph10b subunit, whose C-terminal globular domain matches well with another hook-like density extending towards the adjacent RS2 (Fig. 3F, 1).

Moreover, previous cryo-ET study has demonstrated that in *Tetrahymena*, FAP61 and FAP251 locate in the central stem region and the base of RS3, respectively^43^, we then fitted Cfap61 and Cfap251, the homolog of Fap61 and Fap251, respectively, into similar location in the stem and base of RS3, which matches the structural features well (Fig. 3F, 3). Additionally, Fap91-knockout in *Tetrahymena* resulted in diminished Cfap61 and Cfap251, as well as the RS2-associated Cfap206, suggesting that its homolog Cfap91 might be rooted in the base of RS2 and RS3^39^. We then placed Cfap91 in the corresponding root region of RS3, with its long α-helix extending dramatically upwards to the RS3 head (Fig. 3F)^18,40,43,44^.

Furthermore, our focused 3D classification on the RS1/RS2/RS3 region, respectively, revealed two RS1 conformations relative to the DMT (Fig. 4E), while RS2/RS3 remained relatively stable. Specifically, in one state, RS1 stands more perpendicular to the underneath DMT, while in the other state, RS1 appears more tilted (about 7.6°) towards the adjacent RS3 from the next 96-nm repeat. Distribution analysis showed that the population of the tilted RS1s is approximately 1.6 times of the perpendicular ones, and it appears that the perpendicular RS1s are scattered as small clusters among the tilted RS1s (Fig. 4E). We also observed the occasional “tilt down” of a complete RS1 (Fig. S5C), together indicating a relatively enlarged conformational space of RS1, which may be due to the lacking of constraints from the neighboring RS2/RS3, and that RS1 may be more dynamic in regulating cilia motion.

### Reduced IDAs and MIPs in DMT of mouse ependymal axoneme

In contrast to the helical beat form of sperm flagella, mammalian ependymal and respiratory multicilia beat in a back-and-forth manner but differ greatly in number, length, ciliary beat frequency (CBF), and cellular organization^45–47^. Although cryo-ET maps of mammalian respiratory cilia and sperm flagella are available^16,21,22,41^, to the best of our knowledge, no cryo-ET or cryo-EM structure has been reported for ependymal ciliary axonemes to date. To facilitate a direct comparison among axonemes of different tissues for potential tissue-specific structural variations, we performed further focused refinement on the corresponding regions of the consensus map and acquired *in situ* maps of DMT, IDAs, and the nexin-dynein regulatory complex (N-DRC) in ependymal cilia at a resolution of 19.6 Å for DMT and ∼24.0 Å for the other two (Fig. 4A-D, Table S2).

Strikingly, compared with in situ human respiratory cilia (Fig. 4A) and mouse sperm flagella structures (Fig. S6A), IDA-b/c/e were all missing in ependymal cilia^16^. In addition, IDA-a density is relatively weak (Fig. 4A), and through further 3D classification we found a class with nearly no IDA-a density (Fig. 4F), indicating a variation of IDA-a in different structural repeats. We also managed to fit in AlphaFold2 predicted models of DNAH1, DNAH2, DNAH6, DNAH10, DNAH12, DNAH14, Mbl1, and Mbl2 in the density of IDAs (Fig. 4B), by referring to the homologs in *T. thermophila*^39^ and *Chlamydomonas*^48^. In this proposed model, DNAH12 locates in IDA-a, DNAH1 in IDA-d, DNAH6 in IDA-g, and DNAH10/2 in IDA-I1/fα/fβ (Fig. 4B). Besides, Mbl1 and Mbl2 are located in the stalk of IDA-d as a hetero-dimer (Fig. 4B).

Axonemal microtubule inner proteins (MIPs) associate with the inner surface of the microtubule wall, which are important for microtubule stability and structure. Interestingly, we found that MIPs are also dramatically reduced in mouse ependymal cilia compared with bovine/human respiratory cilia and mouse sperm flagella (Fig. 4C)^16,22,49^. Specifically, Tektin1-4 and Tektin bundle interacting protein 1 (TEKTIP1) within the A-tubule of DMT are all missing in mouse ependymal cilia (Fig. 4C). Similar to ependymal cilia, tektins are also absent in A-tubule of *Chlamydomonas* DMTs (Fig. S6B). In contrast to the mouse ependymal cilia, the mouse sperm flagella exhibit much more filamentous MIPs that is similar to but more extensive than that of tektin filaments inside the A-tubule of DMT (Fig. 4C)^22^. Collectively, these phenomena demonstrate significant tissue specificity for both IDAs and MIPs within DMT, which may be related to the distinct external environment these cilia are facing and also to their distinct pattern of motion.

In addition, the N-DRC in mouse ependymal cilia appears to exhibit a V-shape (Fig. 4D), overall comparable with that in human respiratory cilia^16^, displaying a stronger contact in the upper left corner of the V-shaped density with the B-tubule of the neighboring DMT. We then placed the AlphaFold2 predicted moles of DRC1-5, DRC7-8, and DRC11 subunits into our N-DRC cryo-ET map based on structural features and also by referring to previous cryo-ET study on *Chlamydomonas* N-DRC^48^ and biochemical analysis on mammalian N-DRC^50–53^. In this arrangement, DRC1, DRC2, and two copies of DRC4 assemble into an α-helix bundle going through the entire N-DRC density, DRC5 ties the long α-helix bundle together, DRC8 and DRC11 locate in the upper left position, and DRC3 and DRC7 locate in the middle position, with the extremely long α-helix bundle interacting with almost all DRC subunits (Fig. 4D).

### RS-CP contact patterns

To gain further insights into the RS-CP contact patterns, we incorporated our mouse RS head-neck complex-fitted DMT of mouse ependymal cilia and the cryo-ET map of mouse sperm CP (EMD-27445)^22^ into a “9+2” axoneme by matching remaining RS densities of the CP (Fig. 5A). This fitted multi-scale axoneme structure reveals apparent complementary geometries between the CP projections and the RS heads: (1) CP projection 1d appears to hold the Rsph4a teeth in both RS heads of the DMT8 head-neck dimer, exhibiting a rigid contact mode when they contact during ciliary beat (Fig. 5B-C); (2) CP projections 2b/2d contact the Rsph1 arms from the DMT4 head-neck dimer (Fig. 5B-C), which could lead to elastic RS-CP contact given the intrinsic open-and-close and seesaw-like alternating up-and-down motions happened mainly in the Rsph1 arms of the head-neck dimer (Fig. 2C); (3) CP projection 2d contacts both Rsph1 arms of one RS head and the Rsph4a teeth of the other head of DMT3 (Fig. 5B), exhibiting a combined elastic-and-rigid interaction mode of RS-CP.

Moreover, our analysis of the surface property of the RS head-neck suggested an overall negative electrostatic potential on the CP-facing RS head region, especially for Rsph4a, and a generally hydrophilic property in this region (Fig 5D), collectively indicating a potentially more complex RS-CP interaction mode in mammals^24,25,54^. The complicated geometries and surface properties of projections at and around each presumed contact site might accordingly facilitate the formation of asymmetric planar beat patterns of cilia^10^.

## Discussion

In the present study, we determined the high-resolution cryo-EM structure of the mammalian RS head-neck complex in both monomer and dimer forms (Fig. 1). Our structural analysis revealed a complex subunit interaction network and intrinsic modes of motion of this regulatory machinery of ciliary motility (Fig. 2A, C). We also identified the potential etiology of RS head-neck gene mutations that have been linked to PCD and asthenospermia (Fig. 2B)^13,15,16,42^. Moreover, we present the first *in situ* cryo-ET axoneme structure of mammalian ependymal cilia (Fig. 3), revealing its distinct morphology, especially reduced IDAs and also MIPs within DMT, compared with that of mammalian respiratory cilia and sperm flagella (Fig. 4A-C). In mouse ependymal and mammalian respiratory cilia, multiple sperm-specific densities, including a barrel-shaped density between RS1 and RS2 and densities that cross-link RS2 and RS3, were absent (Fig. S6C)^16,22^. These components are likely trimmed during evolution for their divergent environments and modes of motion. We also built more complete multi-scale models for the RS1/RS2/RS3, IDAs, and N-DRC of mouse ependymal cilia based on our cryo-EM and cryo-ET structures as well as AlphaFold2 predicted models (Fig. 3D-F, 4B, D). Collectively, our study sheds light on the assembly mechanism of the mammalian RS head-neck, the tissue specificity of mammalian muticilia among ependymal and respiratory cilia and flagella, the mode of mammalian RS-CP interaction (Fig. 5A-D), and how these factors collectively regulate ependymal cilia motility.

Based on our cryo-EM studies on the RS head^23^ and head-neck complexes, we propose an assembly mechanism for the mammalian RS head-neck complex (Fig. 5E): (1) the RS head, consisting of two Rsph4a-Rsph9-Rsph1 protomers, can be assembled independently of other subunits; (2) the neck subunit Rsph3b can append to one side of the head and then be wrapped around by the Dpy-30 motif of the head subunits Rsph4a/Rsph4a’ and the neck subunits Rsph2/Rsph23, connecting the head and neck together; (3) Rsph16 may form a dimer independent of other subunits, as it appears as a whole in our RS head-neck monomer structure; (4) the RS head-neck attaches on one side of the arch bridge formed by the Rsph16 dimer, forming the RS head-neck monomer; (5) another head-neck complex can attach to the other side of the arch bridge, forming the RS head-neck dimer for the mammalian RS1/RS2. Through connecting with the extremely long α-helix of Rsph3b, the RS head, neck, and stalk can be assembled into the complete RS1/RS2 complex.

We also proposed a model for mammalian RS3 based on the structural features of the map and model, as well as available biochemical and genetic results (Fig. 3F-G)^18,39,40,43^. The model provides a reliable fit for Cfap251/Cfap61/Cfap91 in the stalk and base region, LRRC23 in the neck, and Rsph9/Rsph4a/Rsph1 in the head of RS3. The fitting of Rsph10b is also reasonable, as its characteristic hook-like feature of the C-terminal globular domain matches the corresponding densities well. In this arrangement, the RS3 head can hold about two RS head complexes, with some extra densities remaining to be filled in. However, the mammalian RS3 head appears slightly larger in size and may have a more complex composition than that of RS1/RS2, indicating a potentially different role of RS3 in RS-CP interaction and cilia motion regulation. We would like to point out that this proposed model for RS3 remains to be validated. A high-resolution structure of mammalian RS3 would be crucial to fully reveal the protein composition and interaction network of its components in atomic detail.

Furthermore, our multi-scale analysis of the RS-CP interaction revealed the rigid contact mode between RS and CP, in addition to the elastic and rigid-elastic RS-CP contact modes (Fig. 5B-C). These rigid RS-CP contacts could constrain the relative movement between CP and RS, which combined with the elastic and rigid-elastic RS-CP contacts could transfer the mechanical force generated by dynein-induced DMT sliding motion^55^ to CP and form constraints between DMT and CP. These factors could potentially mediate the cilia mode of motion, collectively generating an asymmetric planar beat pattern of cilia.

Although flagella and many motile cilia share a common feature of the “9+2” MT arrangement, they exhibit different beating patterns and frequencies^56^. For instance, sperm flagella generally beat in a whip-like motion along a helical path, whereas motile cilia, including tracheal and ependymal cilia, beat in a back-and-forth manner (Fig. 4G). Ependymal cilia are generally longer than tracheal cilia and beat at a faster rate^57^. Related to this, our cryo-ET axoneme structure of mouse ependymal cilia revealed simplified MIPs of DMT without the bundle of tektins in the A-tubule, which differs from that of mammalian respiratory cilia and sperm flagella (Fig. 4C)^22,49^. Since tektins form hyperstable polymers^58^, the absence of tektin bundles within the A-tubule of DMT in ependymal cilia may make them less structurally stiff and stable. Respiratory cilia, ependymal cilia, and sperm flagella function in distinct physiological and environmental contexts. While respiratory cilia and sperm flagella must move the sticky mucus or swim up through the female reproductive tract to reach the egg, ependymal cilia facilitate the circulation of water-like cerebrospinal fluid. Collectively, we propose that ependymal cilia may not require tektins within the A-tubule to provide extra stabilization/stiffness of DMT against waveform-dependent mechanical forces. Substantiating our notion, *Chlamydomonas* flagella, which mostly swim in the water, also lacks the tektin bundle within the A-tubule of its DMT (Fig. S6A)^59^.

The axonemal dynein on DMTs, especially the IDAs which generate the waveform of ciliary/flagellar motion, produces force that is spatiotemporally regulated^60^. Interestingly, our cryo-ET structure revealed a significant reduction in IDAs in ependymal cilia. Specifically, the consecutive IDA-b/c/e in all DMTs are absent, and part of the IDA-a is also missing (Fig. 4A, F). Although it has been reported that IDA-b/e are missing from at least two DMTs in mouse respiratory cilia^61^, and one of the DMTs bears lost IDA-c in *Chlamydomonas*, the complete loss of IDA-b/c/e in all DMTs has not been observed in any other tissues. This indicates that this is a unique ependymal tissue-specific feature. Moreover, it has been reported that a *Chlamydomonas* mutant lacking IDA-c shows only a slight swimming defect in normal culture medium but a greatly reduced swimming ability in viscous media^62^. Therefore, we postulate that IDA-b/c/e may mainly contribute to the force generation in viscous environments. Accordingly, their natural loss in mammalian ependymal cilia is attributed to an adaption to the watery low viscosity CSF during evolution.

It is also interesting to note that, from mammalian sperm flagella to respiratory cilia then to ependymal cilia, ciliary composition and ultrastructure become sequentially simplified. We speculate that this is due to their distinct working environments. For instance, the journey that sperms make through the female reproductive tract to reach an oocyte is long and tortuous. The cervix, for example, contains many folds and grooves filled with mucus^63^. Compared to respiratory cilia that beat in mucus, the increased MIPs and inter-RS connections in sperm flagella (Fig. 4C, S6C)^16,22^ would help to strengthen the stability of sperm flagella and contribute to their whip-like motions. On the other hand, ependymal cilia beat in the water-like environment of brain ventricles. Their reduced IDAs and MIPs (Fig. 4A, C) would thus be accordingly optimized.

In summary, our multi-scale structural study revealed the high-resolution structure of the mammalian RS head-neck complex as well as the unique *in situ* axoneme structure of ependymal cilia. We propose an unprecedented structural model for mammalian RS3, especially for its head, although further examination is needed. Our findings that ependymal cilia lack IDA-b/c/e, have simplified MIPs in DMT, and yet are sufficient to function in a water-like environment highlight an evolutionary choice driven by nature. Our flexibility and multi-scale structural analysis suggested the rigid, elastic, and rigid-elastic RS-CP contacts would constrain the relative movement between CP and RS, transfer the mechanical force between DMT and CP, and mediate the cilia mode of motion to collectively enable an asymmetric planar beat pattern of cilia.

